# Versatile entomopathogenic activity of *Purpureocillium takamizusanense* against diverse agricultural pests

**DOI:** 10.1101/2025.04.08.647886

**Authors:** Pei-Hsin Lo, Khaled Abdrabo El-Sayid Abdrabo, Yu-Shin Nai, Hsiao-ling Lu, Yin-Tse Huang

## Abstract

*Purpureocillium takamizusanense* is an emerging entomopathogenic fungus with significant potential for managing diverse agricultural pests sustainably. This study assesses the broad-spectrum pathogenic activity of *P. takamizusanense* strain TCTeb01, isolated from naturally infected lychee stink bugs (*Tessaratoma papillosa*) in Taiwan. The fungus was evaluated against three economically critical agricultural pests: the coffee berry borer (*Hypothenemus hampei*), southern yellow thrip (*Thrips palmi*), and lychee stink bug. Pathogenicity assays revealed effective control against all tested pests, showing strong pathogenic capabilities and a clear concentration-dependent efficacy. The fungus demonstrated rapid infection and significant pest mortality, highlighting its viability as a biological control agent. Comprehensive morphological and molecular analyses confirmed the fungal strain’s identity and documented additional morphological features, expanding the known variability within the species. Overall, the findings underscore the versatility and robust pathogenic potential of *P. takamizusanense*. Its effectiveness against pests from different insect orders makes it a valuable addition to integrated pest management strategies. Given its environmental compatibility and reduced ecological impact compared to chemical pesticides, *P. takamizusanense* can represent a sustainable alternative for agriculture, contributing positively to pest management practices and ecosystem health.

## Introduction

The genus *Purpureocillium* was established in 2011 to accommodate *Paecilomyces lilacinus* and related species, characterized by their purple to lilac-colored conidia and distinct phylogenetic position within Ophiocordycipitaceae (Luangsa-Ard et al., 2011). Currently, the genus includes eight described species: *P. lilacinum, P. lavendulum, P. takamizusanense, P. atypicola, P. roseum, P. sodanum*, and *P. zongqii*. These fungi exhibit diverse ecological roles, being commonly isolated from soil, decaying vegetation, insects, nematodes, salt crystals, and even as opportunistic pathogens in clinical settings (Calvillo-Medina et al., 2021; Hyde et al., 2016; Luangsa-Ard et al., 2011; Perdomo et al., 2013; Shrestha et al., 2019).

Members of the genus *Purpureocillium* have garnered significant attention as biological control agents due to their broad host range and demonstrated effectiveness against agricultural pests. *Purpureocillium lilacinum*, the most extensively studied species, has been successfully commercialized for controlling plant-parasitic nematodes, particularly *Meloidogyne* species (Dahlin et al., 2019; Singh et al., 2013), which are responsible for substantial annual crop losses worldwide. In addition, *P. lilacinum* has shown efficacy against a variety of pests, including whiteflies (Sani et al., 2023), aphids (Castillo Lopez et al., 2014), spider mites (Silva et al., 2022), and leaf-cutting ants (Goffré and Folgarait, 2015). Similarly, *P. lavendulum* has exhibited effectiveness primarily against nematodes (Bao et al., 2022). The infection mechanisms of these fungi typically involve direct penetration of host cuticles, secretion of extracellular enzymes such as proteases and chitinases, and the production of bioactive metabolites, including leucinostatins, which possess both nematocidal and insecticidal activities (Jiao et al., 2019).

*Purpureocillium takamizusanense*, originally described from cicada adults in Japan (Ban et al., 2015; Kobayashi and Shimizu, 1963), represents an emerging biological control agent with promising potential (Lo and Yu, 2019). Although initial studies focused on its occurrence on cicadas, recent work has documented its natural infection of the lychee stink bug [LSB; *Tessaratoma papillosa* (Drury)] (Lo, 2025; Lo and Yu, 2019), a serious pest of lychee and longan production throughout Southeast Asia that causes significant fruit drop and quality reduction (Yu‐fang and De-xiang, 2000). This finding suggests *P. takamizusanense* may have applications against other economically important insects.

Given the urgent need for sustainable control options against major agricultural pests, *P. takamizusanense* warrants investigation for its efficacy against insects of global economic significance. The coffee berry borer [*Hypothenemus hampei* (Ferrari)] represents coffee’s most destructive pest globally, causing estimated annual losses exceeding $500 million (Durham, 2005). Its cryptic lifecycle inside coffee berries makes conventional control challenging, while its developing resistance to synthetic insecticides further complicates management (Brun et al., 1989; Infante, 2018). Likewise, southern yellow thrip [*Thrips palmi* (Karny)] causes substantial economic damage to vegetable and ornamental crops through both direct feeding damage and transmission of tospoviruses, with global annual losses exceeding $1 billion (MacLeod et al., 2004; Prins and Goldbach, 1998). Despite the economic significance of these pests and the potential of *P. takamizusanense* as a control agent, comprehensive studies on host specificity, infection mechanisms, and efficacy remain limited.

In this study, we provide a detailed characterization of *P. takamizusanense* strains isolated from *T. papillosa*, including expanded morphological descriptions, molecular phylogenetic analyses, and pathogenicity assays against three diverse agricultural pests. Our findings broaden the known host range of *P. takamizusanense* and evaluate its potential as a versatile biological control agent, contributing to the growing body of knowledge on *Purpureocillium* species as important tools in sustainable agriculture.

## Material and Methods

### Fungal isolation

*Purpureocillium takamizusanense* strains were isolated from naturally infected *T. papillosa* collected from longan orchards in Taiwan between 2018 and 2019. Infected insects were examined under a stereomicroscope (Leica DM2000LED) to identify mycosed tissues. Fungal samples were aseptically collected from infected tissues using flame-sterilized scalpels and inoculated onto potato dextrose agar (PDA) plates. The plates were incubated at 25°C and examined for fungal growth at 7-day intervals. Characteristic colonies of *P. takamizusanense* were subcultured onto fresh PDA plates for purification. Pure cultures were maintained on PDA slants at 4°C for experimental use. Reference cultures of all fungal strains were deposited in the PHL’s lab at Taichung District Agricultural Research and Extension Station Bioresource Collection, a reference strain TCTeb01 was deposited at Research Center (BCRC) in Taiwan under accession numbers BCR930208.

### Insect source

We assessed the pathogenicity of *P. takamizusanense* against three agricultural pests: *H. hampei, Thrips palmi*, and *T. papillosa. Hypothenemus hampei* adults were collected from infested *Coffea arabica* berries grown in sun plots in Nantou, Taiwan (23°94’40.9”N 120°95’76.8”E). Adults were obtained using a Mini Insect Breeder (BD7001, MegaView Science Co., Ltd., Taichang, Taiwan) covered with aluminum foil and a strip of kitchen paper in the middle (Wang et al., 2019). Coffee berry borers were reared for more than 14 consecutive generations on an artificial diet (Brun et al., 1993) in glass vials maintained in a growth chamber at 25°C and 85% relative humidity under dark conditions. Female adults of *T. palmi* were sourced from laboratory colonies maintained at the World Vegetable Center. The thrips were reared at 25°C, relative humidity 65-70%, and a photoperiod of 12:12 hr (L:D) in growth chambers. *Tessaratoma papillosa* adults and nymphs were collected from longan orchards in Taiwan. To ensure experimental stability, insects were acclimated on longan branches under controlled conditions (25°C, relative humidity 70%) for three days before experiments.

### Inocula preparation

For all pathogenicity bioassays, we cultured *P. takamizusanense* strain TCTeb01 on potato dextrose agar (PDA) at 25°C for 14 days prior to experimentation. Conidial suspensions were prepared using 0.01% Tween 80 solution. For experiments of *H. hampei* and *T. palmi*, suspensions were adjusted to a concentration of 1×10^8^ conidia/mL, whereas for *T. papillosa* experiments, concentrations of 1×10^5^, 1×10^6^, and 1×10^7^ conidia/mL were used for adult concentration tests and 1×10^7^ conidia/mL for nymph bioassays. Concentration was verified using a hemocytometer. Germination rates for all fungal preparations were confirmed to exceed 90% before use in bioassays by observing spore germination on PDA.

### Pathogenicity test

For bioassay of *H. hampei*, 8 adults per replicate and three replicates were used with each experimental unit. Adults were sprayed with 300 μL of conidial suspension (1×10^8^ conidia/mL) and placed in a Water agar (WA) medium and incubated in a growth chamber at 25 °C in the dark. Control insects were treated with 300 μL 0.01% Tween 80 solution. All boxes were maintained at 25°C in the dark (Chang et al., 2023). Dead insects were removed daily, surface-sterilized with 2% sodium hypochlorite, rinsed twice with sterile water, and placed in moist chambers to confirm mycosis. Survival rate was recorded daily for seven days.

For bioassay of *T. palmi*, four replicates were used with each experimental unit containing ten female adults. The experiment was conducted in plastic cups (55 mm diameter) containing 25 mL of 3% water agar with a circular bean leaf disc (55 mm diameter) on the surface. Each replicate was sprayed with 1 mL of conidial suspension (1×10^8^ conidia/mL) before introduction of thrips. Control groups were treated with 0.01% Tween 80 solution. All cups were covered with 250-mesh netting and lids, then maintained at 25°C with daily monitoring. Dead insects were collected daily, surface-sterilized, and placed in moist chambers to confirm mycosis. Survival rate was recorded daily for seven days.

For bioassay of *T. papillosa*, experimental units contained both nymphs and adults. For nymph tests, insects were sprayed with 3 mL conidial suspension ( 1×10^7^ conidia/mL). For adult tests, three conidial concentrations (1×10^5^, 1×10^6^, and 1×10^7^ conidia/mL) were evaluated. Control insects were treated with 0.01% Tween 80 solution. All insects were maintained at 25°C with 70% relative humidity. Dead insects were collected periodically, surface-sterilized, and placed in moist chambers to confirm mycosis. Survival rate was assessed at days 7, 14, 21, 28, 35, and 42 post-inoculation.

### Fungal morphological and molecular identification

Culture characters were examined on Czapek Dox Agar (CDA; HiMedia Laboratories Pvt. Ltd., India), Malt Extract Agar (MEA; malt extract 20 g, dextrose 20 g, peptone 1 g, agar 20 g, distilled water 1 L), and Potato Dextrose Agar (PDA; HiMedia Laboratories Pvt. Ltd., India). The center-inoculated plates were incubated at 25°C for 7 days. Morphological characters were examined and measured using a Nexcope NE620 compound microscope, and features were documented with a WXCAM microscopic digital camera (Wenxin Instument Co., Ltd., Taiwan). Spores, conidiophores, and conidiogenous apparatus were measured from specimens mounted in 10% potassium hydroxide mount, with at least 50 structures measured for each feature.

For molecular identification, genomic DNA was extracted from the three strains (TCP05, TCTeb01, TP21-2) with our custom Python code. Four gene regions were targeted for phylogenetic analysis: the internal transcribed spacer (ITS), large subunit ribosomal RNA (LSU), RNA polymerase II largest subunit (RPB1), and translation elongation factor 1-alpha (TEF). These regions were identified and extracted from the whole genome sequences of the three strains. Additionally, the ITS, LSU, and TEF regions were amplified from genomic DNA of four other strains (TP17–TP20) using primer sets specific to each targeted region: 983F (GCYCCYGGHCAYCGTGAYTTYAT) and 2218R (ATGACACCRACRGCRACRGTYTG) for TEF (Rehner, 2001), V9G (TACGTCCCTGCCCTTTGTA) and LR1 (GGTTGGTTTCTTTCCT) for ITS (den Ende A. H. G. G. and S., 1999; Vilgalys and Hester, 1990), and LR5 (TCCTGAGGGAAACTTCG) and LROR (ACCCGCTGAACTTAAGC) for LSU (Rehner and Samuels, 1994; Vilgalys and Hester, 1990). For phylogenetic analysis, sequences from related species were retrieved from GenBank (Supplementary Table 1). Multiple sequence alignments were performed using MAFFT v7.490 with default parameters (Katoh and Standley, 2013). The aligned datasets were concatenated into a single dataset after trimming with ClipKIT v2.3.0 using smart-gaps mode to retain phylogenetically informative sites (Steenwyk et al., 2020). ModelTest-NG v.0.1.7 was used to select the best-fit model of DNA evolution using default parameters (Darriba et al., 2020). Maximum likelihood (ML) analyses were conducted using IQ-TREE v2.0.7 with 1000 ultrafast bootstrap replicates (Minh et al., 2020). Bayesian inference was performed using MrBayes v.3.2.7a (Ronquist et al., 2012), with two runs of four chains each, sampling every 100 generations for 1 million generations. *Simplicillium lamellicola* and *S. lanosoniveum* were used as outgroups. Trees were visualized and edited using iTOL (Letunic and Bork, 2024).

### Statistics

Statistical analyses were conducted using R 4.1 with packages including tidyverse (Wickham et al., 2019), car (Fox et al., 2012), lme4 (Bates et al., 2015), emmeans (Lenth et al., 2018), and ggplot2 (Wickham, 2011). For the pathogenicity tests against *H. hampei, T. palmi*, and *T. papillosa* nymphs, one-way Analysis of Variance (ANOVA) was conducted separately at each observation point. F values and corresponding P values were calculated to determine statistical significance of treatment effects at each time point.

For the concentration-dependent experiment on adult *T. papillosa*, a mixed model ANOVA was employed to analyze the effects of concentration (10^5^, 10^6^, and 10^7^ conidia/mL), days post-inoculation, and their interaction on mortality rates. The statistical model included concentration and time as fixed effects, with replicates as random effects. Post-hoc comparisons between different concentrations at each observation point (days 7, 14, 21, 28, 35, and 42) were conducted using Tukey’s HSD test to determine significant differences between specific treatment pairs.

Median lethal time (LT_50_) values and their 95% confidence intervals were calculated using dose-response modeling with the drc package (Ritz and Streibig, 2005). Time-mortality data were fitted to log-logistic models (LL.2 or LL.3) based on model convergence criteria. For treatments where log-logistic models did not adequately fit, probit regression models were applied.

## Results

### Pathogenicity test

#### Pathogenicity against agricultural pests

*Purpureocillium takamizusanense* strain TCTeb01 demonstrated significant pathogenicity against all three agricultural pests tested (Figure 1, Table 1). The fungus led to complete reduction in survival (0%) of *H. hampei* by day 7, reduced *T. palmi* survival to 22.5% by day 7, and decreased *T. papillosa* nymph survival to 32.5% after 42 days.

**Table 1.** Survival rates of three agricultural pests inoculated with *P. takamizusanense* TCTeb01 at different observation points. Significant differences compared to the control group, assessing using ANOVA, are indicated.

| Observation point † | Pest species | Control survival (%) | Treatment survival (%) | F value | P value |
| --- | --- | --- | --- | --- | --- |
| 1 | *H. hampei* | 95.83 | 100 | 1 | 0.374 |
|  | *T. palmi* | 100 | 97.5 | 1 | 0.356 |
|  | *T. papillosa nymph* | 97.5 | 86.25 | 5.4 | 0.049* |
| 2 | *H. hampei* | 95.83 | 91.67 | 0.2 | 0.678 |
|  | *T. palmi* | 92.5 | 92.5 | 0 | 1.000 |
|  | *T. papillosa nymph* | 92.5 | 70.62 | 2.12 | 0.183 |
| 3 | *H. hampei* | 87.5 | 70.83 | 0.84 | 0.411 |
|  | *T. palmi* | 90 | 70 | 3 | 0.134 |
|  | *T. papillosa nymph* | 90 | 55.62 | 5.56 | 0.046* |
| 4 | *H. hampei* | 83.33 | 33.33 | 4.97 | 0.090 |
|  | *T. palmi* | 85 | 50 | 9.8 | 0.020* |
|  | *T. papillosa nymph* | 82.5 | 40.62 | 9.83 | 0.014* |
| 5 | *H. hampei* | 83.33 | 16.67 | 12.8 | 0.023* |
|  | *T. palmi* | 80 | 40 | 24 | 0.003** |
|  | *T. papillosa nymph* | 82.5 | 34.37 | 12.98 | 0.007** |
| 6 | *H. hampei* | 79.17 | 0 | 361 | 4.52E-05** |
|  | *T. palmi* | 77.5 | 30 | 40.11 | 0.001** |
|  | *T. papillosa nymph* | 80 | 32.5 | 13.13 | 0.007** |
| 7 | *H. hampei* | 70.83 | 0 | 289 | 7.02E-05** |
|  | *T. palmi* | 77.5 | 22.5 | 66 | 1.87E-04** |

† *H. hampei* and *T. palmi*: observation points 1-7 = days 1-7
† *T. papillosa nymph*: observation points 1-6 = days 7, 14, 21, 28, 35, and 42

**Figure 1.**
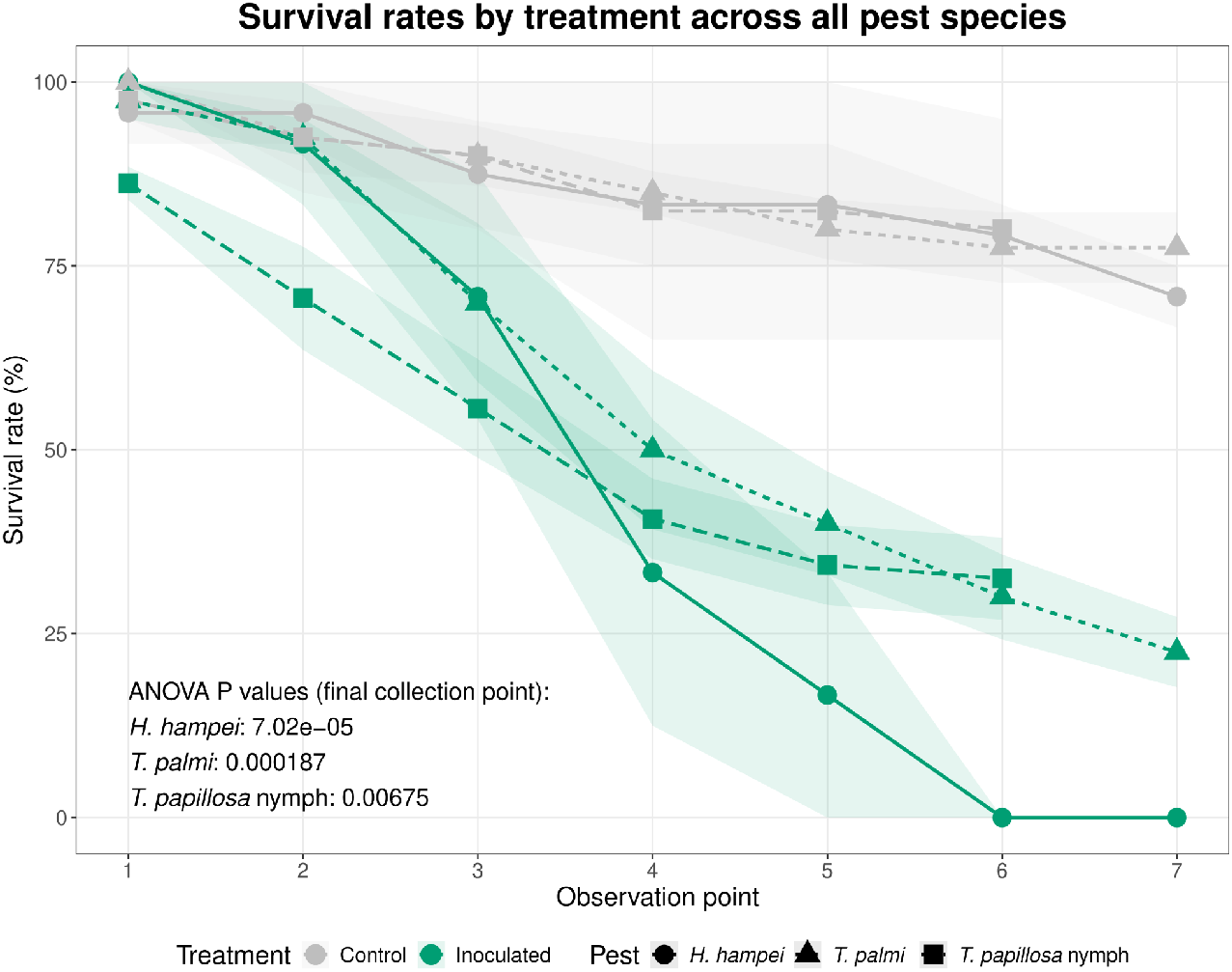
Survival of three agricultural pests treated with *P. takamizusanense* TCTeb01. Survival rates were significantly lower in inoculated treatments (green) compared to controls (gray) across all pest species. Shade areas represent standard error. Observation point 1-7 for *H. hampei* and *T. palmi* are 1-7 days after inoculation (DPI); for *T. papillosa* nymph, they are 7, 14, 21, 28, 35, and 42 DPI.

Survival of *H. hampei* declined rapidly from 100% on day 1 to 70.83% by day 3, dropped further to 33.33% by day 4, and reached 0% by days 6–7. *Thrips palmi* displayed a steady decrease in survival, from 97.5% at day 1 down to 70% by day 3, further declining to 50% by day 4, and finally reaching 22.5% by day 7. *Tessaratoma papillosa* nymphs exhibited a more moderate decline, with survival at 86.25% on day 7, gradually decreasing to 55.62% by day 21, and ultimately dropping to 32.5% by day 42 (Figure 1, Table 1).

Significant differences in survival for *H. hampei* became evident from day 5 and were highly significant by days 6–7 (Table 1). For *T. palmi*, statistical significance in survival rates emerged by day 4 and strengthened in subsequent days (Table 1). In *T. papillosa* nymphs, significant reductions in survival were observed at days 7, 21, 28, 35, and 42, with the most substantial statistical differences recorded at the final observation point (Table 1).

#### Pathogenicity against adult *T. papillosa* at different concentrations

Three conidial concentrations (10^5^, 10^6^, and 10^7^ conidia/mL) demonstrated a concentration-dependent effect on adult mortality, with the highest concentration (10^7^ conidia/mL) producing greater mortality rates throughout the experimental period (Figure 2). By day 42, mortality rates reached 90.0% at 10^7^ conidia/mL, compared to 59.0% and 37.0% at 10^5^ and 10^6^ conidia/mL, respectively. The mortality rates differed significantly between concentrations, with distinct temporal patterns observed across treatments. Mixed model ANOVA confirmed significant effects of concentration (F = 4.86, df = 2, 46, P = 0.012), days post-inoculation (F = 176.08, df = 1, 46, P = 2.45E-017), and their interaction (F = 5.56, df = 2, 46, P = 0.007) on adult *T. papillosa* mortality. These results indicate that both factors significantly influenced insect mortality

**Figure 2.**
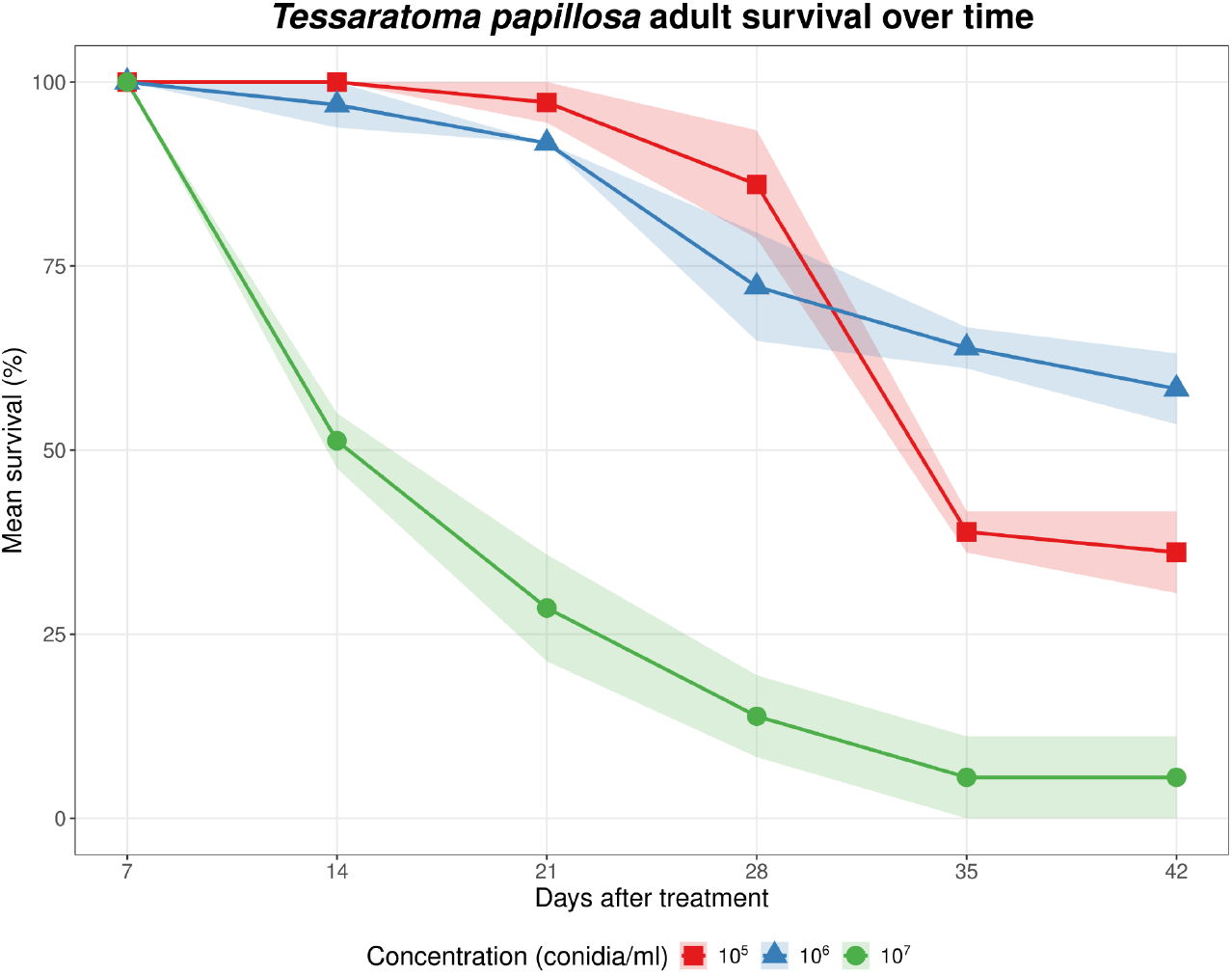
Effect of *P. takamizusanense* TCTeb01 conidial concentration on survival of adult *T. papillosa*. Three concentrations (10^5^, 10^6^, and 10^7^ conidia/mL) were tested over 42 days. The 10^7^ conidia/mL demonstrated the greatest reduction in survival, with only 5% of adults surviving by day 35. Shaded areas represent 95% confidence intervals.

No mortality was observed across treatment groups during the first 7 days post-inoculation. The highest concentration (10^7^ conidia/mL) began showing efficacy by day 14 (50.0% mortality), significantly exceeding the lower concentrations. From days 21–28, the 10^7^ treatment maintained significantly higher mortality compared to both 10^5^ and 10^6^ treatments. At day 35, all three concentrations differed significantly from each other, with the 10^5^ concentration (52.0%) exceeding the 10^6^ concentration. By day 42, the 10^7^ concentration reached 90.0% mortality, significantly higher than both lower concentrations. The concentration-dependent mortality pattern indicated that 10^7^ conidia/mL was the most effective treatment against adult *T. papillosa* (Table 2).

**Table 2.** Pairwise comparisons of survival rates in adult *T. papillosa* exposed to varying concentrations of *P. takamizusanense* TCTeb01. Significant differences between treatments are indicated.

| DPI<sup>†</sup> | Comparison | Estimate | SE | df | t-ratio | P value |
| --- | --- | --- | --- | --- | --- | --- |
| 7 | 10<sup>5</sup> vs 10<sup>6</sup> | 0 | 0 | 6 | NA | NA |
|  | 10<sup>5</sup> vs 10<sup>7</sup> | 0 | 0 | 6 | NA | NA |
|  | 10<sup>6</sup> vs 10<sup>7</sup> | 0 | 0 | 6 | NA | NA |
| 14 | 10<sup>5</sup> vs 10<sup>6</sup> | -3.1 | 3.97 | 6 | -0.78 | 0.728 |
|  | 10<sup>5</sup> vs 10<sup>7</sup> | -48.73 | 3.97 | 6 | -12.269 | 4.40E-05** |
|  | 10<sup>6</sup> vs 10<sup>7</sup> | -45.63 | 3.97 | 6 | -11.489 | 6.45E-05** |
| 21 | 10<sup>5</sup> vs 10<sup>6</sup> | -5.53 | 6.33 | 6 | -0.874 | 0.675 |
|  | 10<sup>5</sup> vs 10<sup>7</sup> | -68.67 | 6.33 | 6 | -10.845 | 8.98E-05** |
|  | 10<sup>6</sup> vs 10<sup>7</sup> | -63.13 | 6.33 | 6 | -9.971 | 1.45E-04** |
| 28 | 10<sup>5</sup> vs 10<sup>6</sup> | -13.9 | 9.63 | 6 | -1.444 | 0.379 |
|  | 10<sup>5</sup> vs 10<sup>7</sup> | -72.23 | 9.63 | 6 | -7.502 | 0.001* |
|  | 10<sup>6</sup> vs 10<sup>7</sup> | -58.33 | 9.63 | 6 | -6.058 | 0.002* |
| 35 | 10<sup>5</sup> vs 10<sup>6</sup> | 25 | 5.58 | 6 | 4.482 | 0.010* |
|  | 10<sup>5</sup> vs 10<sup>7</sup> | -33.33 | 5.58 | 6 | -5.976 | 0.002* |
|  | 10<sup>6</sup> vs 10<sup>7</sup> | -58.33 | 5.58 | 6 | -10.458 | 1.10E-04** |
| 42 | 10<sup>5</sup> vs 10<sup>6</sup> | 22.2 | 7.54 | 6 | 2.945 | 0.058 |
|  | 10<sup>5</sup> vs 10<sup>7</sup> | -30.57 | 7.54 | 6 | -4.055 | 0.016* |
|  | 10<sup>6</sup> vs 10<sup>7</sup> | -52.77 | 7.54 | 6 | -7.001 | 0.001* |

<sup>†</sup> Days post-inoculation

#### Lethal time analysis across pest species

The lethal time to 50% mortality (LT_50_) varied significantly among pest species and conidial concentrations (Table 3). For adult *T. papillosa*, LT_50_ values demonstrated a concentration-dependent response, with the highest concentration (1×10^7^ conidia/mL) achieving LT_50_ in 17.8 days (95% CI: 9.9-25.7), substantially faster than the 36.3 days (CI: 28.5-44.2) and 42.3 days (CI: 23.7-60.8) observed at 1×10^5^ and 1×10^6^ conidia/mL, respectively. Nymphs of *T. papillosa* showed comparable susceptibility to adults at the highest concentration, with an LT_50_ of 20.1 days (CI: 4.2-35.9). Both *H. hampei* and *T. palmi* exhibited remark susceptibility, with LT_50_ values of 3.6 days (CI: 2.8-4.5) and 3.7 days (CI: 2.5-4.9), respectively, when treated with 1×10^8^ conidia/mL.

**Table 3.** LT_50_ of insect pest species inoculated by *P. takamizusanense* TCTeb01 at varied concentrations

| Pest | Life stage | Inoculum<br>concentration conidia<br>ml<sup>-1b</sup> | LT<sub>50</sub> (days) | Lower CI | Upper CI |
| --- | --- | --- | --- | --- | --- |
| *T. papillosa* | Adult | 1×10<sup>5</sup> | 36.3 | 28.5 | 44.2 |
| *T. papillosa* | Adult | 1×10<sup>6</sup> | 42.3 | 23.7 | 60.8 |
| *T. papillosa* | Adult | 1×10<sup>7</sup> | 17.8 | 9.9 | 25.7 |
| *T. papillosa* | nymph | 1×10<sup>7</sup> | 20.1 | 4.2 | 35.9 |
| *H. hampei* | Adult | 1×10<sup>8</sup> | 3.6 | 2.8 | 4.5 |
| *T. palmi* | Adult | 1×10<sup>8</sup> | 3.7 | 2.5 | 4.9 |

### Fungal morphological and molecular identification

#### Morphological characterization

The morphological characteristics of strains TCTeb01 was examined on various culture media (Figure 3A-C). Colonies appeared floccose to powdery in texture, pale vinaceous to lilac coloration at the center and whitish toward the margins. The reverse was distinctly yellowish to light brown, showing a diffusible pigment.

**Figure 3.**
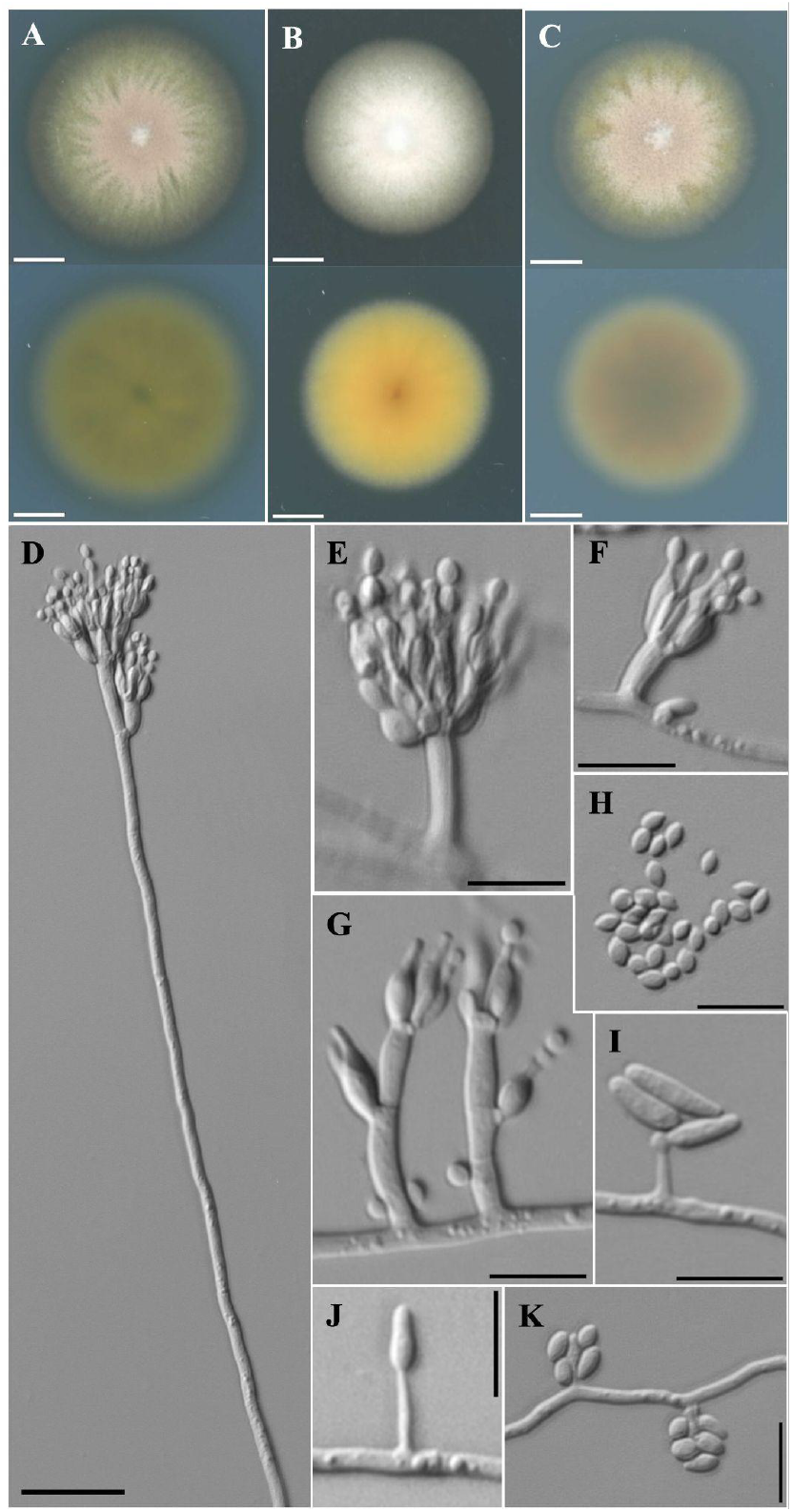
*Purpureocillium takamizusanensis* (TCTeb01). A–C. Colonies at 25 C after 7 d. A. Czapek Dox Agar. B. Malt Extract Agar. C. Potato Dextrose Agar. D. Conidiophores arising from aerial hyphae with biverticillate penicillus growth. E–G. Conidiophores arising from effuse hyphae with monoverticillate penicillus growth. H. Smooth-walled and ellipsoidal conidia. I–K. *Acremonium*-like synanamorph with typical cylindrical (I–J), and ellipsoidal conidia in slimy masses (K). Scale bars; A–C = 5 mm, D = 20 μm, E–K = 10 μm.

*Purpureocillium takamizusanense* (Kobayasi) Ban, Azuma & Hirok. Sato Conidiophores dimorphic: Long erect conidiophores 94-169 μm × 2.5-3 μm, smooth to rough-walled; Short irregularly branched conidiophores 5.7-9.9 × 3.1-3.2 μm. Phialides elongated clavate or cylindrical with tapered apex, 9.2-11 × 2-2.1 μm. Conidia in basipetal chains, smooth-walled, ellipsoidal, 2-2.9 × 1.5-1.9 μm. Acremonium-like synanamorph consistently present in all examined strains with: Conidiophores 5.3-7.1 μm long; Two types of conidia - (1) Cylindrical slightly curved conidia 6.3-8.7 × 1.8-1.9 μm, and (2) Ellipsoidal conidia in slimy masses 2.3-3.2 × 1.6-2 μm (Figure 3).

The Taiwanese strains show similar overall morphology to the type specimen from Japan (Ban et al., 2015; Kobayashi and Shimizu, 1963), which was described as having smaller conidiophores (65-110 μm long) and slightly larger conidia (2.5-4 × 1.4-1.8 μm). Our observations extend the known morphological variation of this species, particularly in conidiophore length (up to 169 μm) and document the consistent presence of an Acremonium-like synanamorph with two distinct conidial types. This synanamorph feature was not explicitly described in the original Japanese specimens.

### Molecular phylogenetic analysis

To determine the taxonomic placement of the seven strains, phylogenetic analyses were performed using a concatenated dataset comprising four genetic loci: ITS, LSU, TEF, and RPB1, resulting in a total alignment length of 4126 bp. The best-fit substitution models, selected by ModelTest-NG, were TrN for ITS, TIM2+G4 for LSU, K80+G4 for RPB1, and K80+I+G4 for TEF.

The maximum likelihood phylogeny (Figure 4) clearly demonstrated that our seven Taiwanese isolates (TCP05, TCTeb01, TP21-2, TP17, TP18, TP19, and TP20) formed a strongly supported monophyletic group within the species *P. takamizusanense*. These isolates clustered closely together and subsequently formed a robustly supported larger subclade with the syntype strains of *P. takamizusanens*e, NBRC 108982 and NBRC 110232, representing partspore and conidial isolates from the same specimen. This subclade was sister to another *P. takamizusanense* group, including NBRC 100742 and strain F1724, which was supported with moderate confidence.

**Figure 4.**
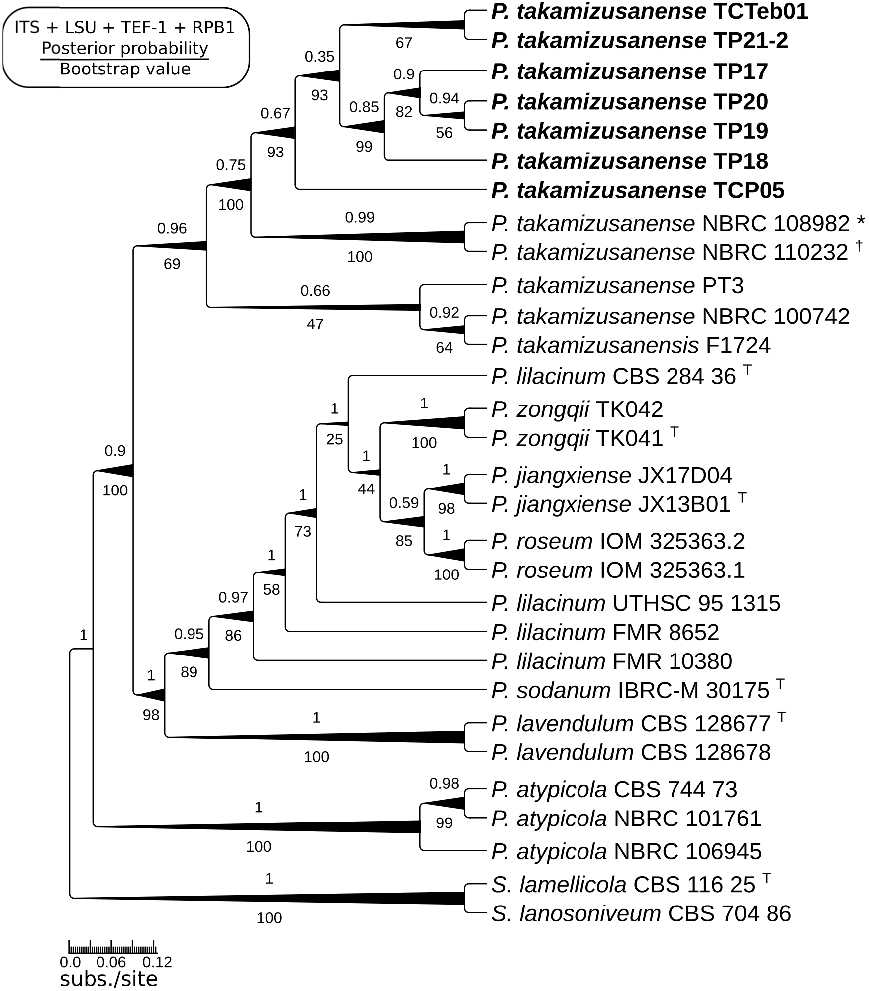
Phylogeny obtained by Maximum likelihood estimation of the combined ITS + LSU + TEF-1 + RPB1 sequences data showing the phylogenetic relationships within the genus *Purpureocillium. Simplicillium lamellicola* and *S. lanosoniveum* were selected as outgroups. Numbers above nodes represent Bayesian posterior probability values; numbers below nodes represent maximum likelihood bootstrap values. Branch widths are proportional to bootstrap values and posterior probability values. Type strains are indicated with superscript T. Asterisk (*) indicates strain derived from partspores, and dagger (†) indicates strain derived from conidia according to Perdomo et al. (2013). Scale bar indicates 0.06 substitutions per site.

Our multilocus phylogeny distinguished the *P. takamizusanense* clade from other *Purpureocillium* species, with most species-level relationships receiving high bootstrap values (>90%) and high posterior probabilities (>0.95). Bayesian inference yielded a congruent topology that strongly supported the same evolutionary relationships observed in the maximum likelihood analysis.

## Discussion

This study provides the first comprehensive characterization of *P. takamizusanense* as a versatile entomopathogenic fungus with significant activity against multiple agricultural pests. Our results demonstrate that *P. takamizusanense* strain TCTeb01 exhibits strong pathogenicity against coffee berry borer (*H. hampei*), southern yellow thrip (*Thrips palmi*), and lychee stink bug (*T. papillosa*), expanding its known host range beyond the originally described cicada hosts.

### Comparative pathogenicity and host range expansion

Our findings demonstrate that *P. takamizusanense* strain TCTeb01 exhibits significant pathogenicity against three economically important pests from different insect orders. When compared with other entomopathogenic fungi tested against *H. hampei*, our strain achieved complete mortality (100%) by day 7 with an LT_50_ of 3.6 days (Table 3). This efficacy is comparable to several *Beauveria bassiana* strains in the previous studies. For example, *B. bassiana* strains CCB-LE265 and P19 showed similar LT_50_ values ranging from 2.52 to 2.9 days at equivalent or higher concentrations (Chuquibala-Checan et al., 2023). However, our strain outperformed other reported agents such as *B. bassiana* strain CG11 (LT_50_ = 7 days) and *Metarhizium anisopliae* strain CG46 (LT_50_ = 9.4 days)(Samuels et al., 2002).

For *T. palmi, P. takamizusanense* TCTeb01 achieved 77.5% mortality by day 7 with an LT_50_ of 3.7 days at 1×10^8^ conidia/mL. This performance is comparable to *B. bassiana* strain JEF-350 (LT_50_ = 3.73 days) and slightly less effective than *M. lepidiotae* strain NCHU-9 (LT_50_ = 2.89 days) at the same concentration (Li et al., 2021; Mushyakhwo et al., 2025). The mortality rate achieved by our strain suggests it could serve as an effective biological control alternative for this challenging pest.

Against *T. papillosa*, our strain showed a concentration-dependent efficacy pattern. At 1×10^7^ conidia/mL, *P. takamizusanense* achieved 94.4% mortality of adults by day 42 with an LT_50_ of 17.8 days. While this represents a longer time to mortality compared to *B. bassiana* strains tested against *T. papillosa* (LT_50_ values ranging from 5.2 to 7.85 days) (Lin, 2005; Meng et al., 2017; Su, 2021), our strain eventually achieved higher overall mortality rates. Importantly, *P. takamizusanense* outperformed *P. lilacinus* strain Ta-01 against adult *T. papillosa*, which had an LT_50_ of 8.35 days but with lower ultimate mortality (63.33% by day 10) (Meng et al., 2017).

The comparative efficacy data suggests that while *P. takamizusanense* may not act as rapidly as some highly virulent *B. bassiana* strains against certain pests, it exhibits a broader host range and ultimately achieves high mortality rates. This versatility across pest species from different insect orders represents a valuable attribute for integrated pest management strategies.

### Host diversity and biosafety profile

The expanded host range of *P. takamizusanense* documented in this study enhances its potential utility as a versatile biological control agent. The effectiveness against pests from three distinct insect orders—Coleoptera (*H. hampei*), Thysanoptera (*T. palmi*), and Hemiptera (*T. papillosa*)—demonstrates a broader spectrum of activity than previously known for this species. This cross-order efficacy is particularly valuable for agricultural systems where multiple pest groups co-occur.

The variable LT_50_ values observed across pest species suggest that host-specific factors influence the infection process and mortality rate. The faster mortality in smaller insects (*H. hampei* and *T. palmi*) compared to the larger stink bug (*T. papillosa*) may reflect differences in cuticle thickness, immune responses, or other physiological barriers that affect fungal penetration and proliferation (Bitencourt et al., 2021; Mannino et al., 2019; Ortiz-Urquiza and Keyhani, 2013; Ramirez et al., 2018).

Within the genus *Purpureocillium, P. lilacinum* has been extensively studied for its efficacy against plant-parasitic nematodes (Dahlin et al., 2019; El-Marzoky et al., 2023; Khan and Tanaka, 2023), whiteflies (Sani et al., 2023), aphids (Castillo Lopez et al., 2014), and spider mites (Silva et al., 2022), with several commercial formulations available. In contrast, this study represents the first comprehensive evaluation of *P. takamizusanense* as a biological control agent beyond its natural association with cicadas and lychee stink bugs. The demonstrated broad host spectrum across multiple insect orders suggests that *P. takamizusanense* possesses substantial potential for integrated pest management programs. Additionally, preliminary biosafety assessments indicate this fungus has low pathogenicity to non-target organisms, including beneficial insects and aquatic species, further enhancing its appeal as a sustainable alternative to chemical pesticides for controlling diverse agricultural pests (Lo, 2025).

### Morphological characterization and taxonomic significance

Morphologically, our Taiwanese strains of *P. takamizusanense* align with the type specimen from Japan, while extending the known morphological variation of this species. The conidiophore length observed (up to 169 μm) exceeds previously reported dimensions (65-110 μm), and we consistently documented an *Acremonium*-like synanamorph with two distinct conidial types. This synanamorph feature was not explicitly described in the original Japanese specimens (Ban et al., 2015; Kobayashi and Shimizu, 1963), suggesting either morphological plasticity or regional variation within the species. The distinct yellowish to light brown diffusible pigment on culture media further distinguishes these strains and may have taxonomic significance.

## Conclusions

This study demonstrates that *P. takamizusanense* possesses significant potential as a versatile biological control agent against diverse agricultural pests. Its ability to infect and kill insects from different orders (Coleoptera, Thysanoptera, and Hemiptera) represents a valuable attribute for integrated pest management strategies. The expanded morphological description and molecular characterization provided here contribute to the taxonomic understanding of this species and facilitate its accurate identification in future studies.

Further research should focus on optimizing application methods, assessing field efficacy, infection mechanisms, and investigating potential synergistic effects with other control measures to fully exploit the biocontrol potential of *P. takamizusanense* in sustainable agriculture.

## Acknowledgement

We sincerely thank Mr. Chauang, Hoang-More, the owner of the coffee orchard in Nantou, for providing the dry, infested coffee berries.

## Legend

**Supplementary Table 1.** Strain source information, NCBI accession numbers, and references for *Purpureocillium* species used in the phylogenetic analysis.

| Species | Strain | Source | ITS | LSU | RPB1 | TEF | Reference | Note |
| --- | --- | --- | --- | --- | --- | --- | --- | --- |
| *P. sodanum* | IBRC-M 30175 T | Salt crystals | KX668542 | - | - | - | Hyde et al. 2016 |  |
| *P. lilacinum* | FMR 8652 | Human | FR734090 | FR775473 | - | - | Perdomo et al. 2013 |  |
| *P. lavendulum* | CBS 128678 | Human | MH864977 | MH876430 | - | - | Perdomo et al. 2013 |  |
| *P. roseum* | IOM 325363.1 | Human | MT560195 | MT560197 | - | - | Calvillo-Medina et al. 2021 |  |
| *P. roseum* | IOM 325363.2 | Human | MT560196 | MT560198 | - | - | Calvillo-Medina et al. 2021 |  |
| *P. atypicola* | CBS 744.73 | Atypus karschi | GU980041 | EF468841 | EF468786 | - | Perdomo et al. 2013 |  |
| *P. lilacinum* | CBS 431.87 | *Meloidogyne* eggs, Philippines | AY624188 | EF468844 | EF468897 | EF468791 | Perdomo et al. 2013 |  |
| *P. lilacinum* | FMR 8250 | Nematode eggs, Amposta, Spain | FR734086 | FR775469 | FR775492 | FR734141 | Perdomo et al. 2013 |  |
| *P. lilacinum* | FMR 10040 | Soil, Ira | FR734092 | FR775475 | FR775498 | FR734147 | Perdomo et al. 2013 |  |
| *P. lilacinum* | FMR 10041 | Soil, Cuba | FR734093 | FR775476 | FR775499 | FR734148 | Perdomo et al. 2013 |  |
| *P. lilacinum* | FMR 10068 | Soil, Nsukka, Nigeria | FR734094 | FR775477 | FR775500 | FR734149 | Perdomo et al. 2013 |  |
| *P. lilacinum* | CBS 284.36 T | Soil | FR734101 | FR775484 | FR775507 | FR734156 | Perdomo et al. 2013 |  |
| *P. lilacinum* | JCM 8437 | Heterodera zeae, Kent County, USA | FR734103 | FR775486 | FR775509 | FR734158 | Perdomo et al. 2013 | Paecilomyces nostocoides T |
| *P. lilacinum* | JCM 8372 | Litter, Sado Island, Japan | FR734104 | FR775487 | FR775510 | FR775514 | Perdomo et al. 2013 |  |
| *P. lilacinum* | FMR 10380 | Soil, Reus, Spain | FR734102 | FR775485 | FR775508 | FR734157 | Perdomo et al. 2013 |  |
| *P. lilacinum* | UTHSC 95-13 | Scalp, USA | FR734099 | FR775482 | FR775505 | FR734154 | Perdomo et al. 2013 |  |
| *P. lilacinum* | JCM 8438 | Heterodera zeae, Kent County, USA | FR734105 | FR775488 | FR775511 | FR775515 | Perdomo et al. 2013 | Paecilomyces nostocoides |
| *P. lavendulum* | CBS 128677 T | Soil | FR734106 | - | - | FR775516 | Perdomo et al. 2013 |  |
| *P. takamizusanensis* | F1724 | Unknown | GU980039 | - | - | GU979993 | Perdomo et al. 2013 | Isaria takamizusanensis |
| *P. takamizusanense* | NBRC 100742 | Tanna japonensis | LC008197 | - | - | LC008333 | Ban et al. 2015 |  |
| *P. takamizusanense* | NBRC 108982 | Cicada | LC008204 | - | - | LC008338 | Ban et al. 2015 |  |
| *P. takamizusanense* | NBRC 110232 | - | LC008205 | - | - | LC008339 | Ban et al. 2015 |  |
| *P. takamizusanense* | TCP05 | *Tessaratoma papillosa* | PV382381 | PV382390 | PV394270 | PV394266 | This sutdy | Extracted from genome |
| *P. takamizusanense* | TCTeb01 | *Tessaratoma papillosa* | PV382380 | PV382389 | PV394269 | PV394265 | This sutdy | Extracted from genome |
| *P. takamizusanense* | TP21-2 | *Tessaratoma papillosa* | PV382379 | PV382388 | PV394268 | PV394264 | This sutdy | Extracted from genome |
| *P. takamizusanense* | TP17 | *Tessaratoma papillosa* | PV382383 | PV382392 | - | PV394261 | This sutdy |  |
| *P. takamizusanense* | TP18 | *Tessaratoma papillosa* | PV382384 | PV382393 | - | PV394267 | This sutdy |  |
| *P. takamizusanense* | TP19 | *Tessaratoma papillosa* | PV382385 | PV382394 | - | PV394263 | This sutdy |  |
| *P. takamizusanense* | TP20 | *Tessaratoma papillosa* | PV382386 | PV382395 | - | PV394262 | This sutdy |  |
| *P. takamizusanense* | PT3 | *Meimuna opalifera* | PV382382 | PV382391 | PV394271 | PV394260 | This sutdy | Extracted seqs from GCA 022605165.1 |
| *P. atypicola* | NBRC 106945 | Spider, Spinosa sp., Okinawa | LC008212 | CC01332301 | - | LC008346 | Ban et al. 2015 |  |
| *P. atypicola* | NBRC 101761 | Spider, Osaka | LC008213 | CC00823501 | - | LC008347 | Ban et al. 2015 |  |
| *P. jiangxiense* | JX17D04 | Soil | PP555636 | PP555645 | - | PP658209 | Chen et al. 2024 |  |
| *P. jiangxiense* | JX13B01 T | Soil | PP555637 | PP555646 | - | PP658210 | Chen et al. 2024 |  |
| *P. zongqii* | TK041 T | Soil | PQ211280 | PQ211284 | - | PQ223681 | Chen et al. 2024 |  |
| *P. zongqii* | TK042 | Soil | PQ211281 | PQ211285 | - | PQ223682 | Chen et al. 2024 |  |
| *Simplicillium lamellicola* | CBS 116.25 T | *Agaricus bisporus* | MH854806 | AF339552 | DQ522462 | DQ522356 | Luangsa-ard et al. 2011 | outgroup |
| *S. lanosoniveum* | CBS 704.86 | *Hemileia vastatrix* | AJ292396 | AF339553 | DQ522464 | DQ522358 | Luangsa-ard et al. 2011 | outgroup |

